# GCNG: Graph convolutional networks for inferring cell-cell interactions

**DOI:** 10.1101/2019.12.23.887133

**Authors:** Ye Yuan, Ziv Bar-Joseph

**Affiliations:** Machine Learning Department, School of Computer Science, Carnegie Mellon University, Pittsburgh, PA 15213, USA; Computational Biology Department, School of Computer Science, Carnegie Mellon University, Pittsburgh, PA 15213, USA

**Keywords:** Graph convolutional network, Spatial transcriptomics, Gene relationship inference

## Abstract

Several methods have been developed for inferring gene-gene interactions from expression data. To date, these methods mainly focused on intra-cellular interactions. The availability of high throughput spatial expression data opens the door to methods that can infer such interactions both within and between cells. However, the spatial data also raises several new challenges. These include issues related to the sparse, noisy expression vectors for each cell, the fact that several different cell types are often profiled, the definition of a neighborhood of cell and the relatively small number of extracellular interactions. To enable the identification of gene interactions between cells we extended a Graph Convolutional Neural network approach for Genes (GCNG). We encode the spatial information as a graph and use the network to combine it with the expression data using supervised training. Testing GCNG on spatial transcriptomics data we show that it improves upon prior methods suggested for this task and can propose novel pairs of extracellular interacting genes. Finally, we show that the output of GCNG can also be used for down-stream analysis including functional assignment.

Supporting website with software and data: https://github.com/xiaoyeye/GCNG.

## 1 Introduction

Several computational methods have been developed over the last two decades to infer interaction between genes based on their expression [22]. Early work utilized large compendiums of microarray data [28] while more recent work focused on RNA-Seq and scRNA-Seq [26]. While the identification of pairwise interactions was the goal of several studies that relied on such methods, others used the results as features in a classification framework [15] or as pre-processing steps for the reconstruction of biological interaction networks [4]. Most work to date focused on intra-cellular interactions and network. In such studies we are looking for interacting genes involved in a pathway or in the regulation of other genes within a specific cell. In contrast, studies of extracellular interactions (i.e. interactions of genes or proteins in different cells) mainly utilized small scale experiments in which a number of ligand and receptors pairs were studied in the context of a cell line or tissue [20]. However, recently developed methods for spatial transcriptomics are now providing high throughput information about both, the expression of genes within a single cell and the spatial relationships between cells [5, 14, 17, 23, 27]. Such information opens the door to much larger scale analysis of extracellular interactions.

Current methods for inferring extracellular interactions from spatial transcriptomics have mostly focused on unsupervised correlation based analysis. For example, Giotto method calculated effect upon gene expression from neighbor cell types [7]. While these approaches perform well in some cases, they may not identify interactions that are limited to a specific area, specific cell types or that are related to more complex patterns (for example, three way interactions).

To overcome these issues we present a new method that is based on graph convolutional neural networks (GCNs). GCNs have been introduced in the machine learning literature a few years ago [2]. Their main advantage is that they can utilize the power of convolutional NN even for cases where spatial relationships are not complete [29, 31]. Specifically, rather than encoding the data using a 2D matrix (or a 1D vector) GCNs use the graph structure to encode relationships between samples. The graph structure (represented as a normalized interaction matrix) is deconvolved together with the information for each of the nodes in the graph leading to NN that can utilize both, the values encoded in each node (in our case gene expression) and the relationship between the cells expressing these genes.

To apply GCN to the task of predicting extracellular interactions from gene expression (GCNG) we first convert the spatial transcriptomics data to a graph representing the relationship between cells. Next, for each pair of genes we encode their expression and use GCNG to convolve the graph data with the expression data. This way the NN can utilize not just first order relationships, but also higher order relationships in the graph structure. We discuss the specific transformation required to encode the graph and gene expression, how to learn parameters for the GCNG and how to use it to predict new interactions.

We tested our approach on data from the two spatial transcriptomics methods that profile the most number of genes right now, SeqFISH+ [8] and MERFISH [30]. As we show, GCNG greatly improves upon correlation based methods when trying to infer extracellular interactions. We visually analyze some of the correctly predicted pairs and show that GCNG can overcome some of the limitations of unsupervised methods by focusing on only a relevant subset of the data. Analysis of the predicted pairs shows that many are either known to interact or know to be involved in a similar functional pathway supporting their top ranking.

## 2 Methods

We extended ideas from GCN and developed a general supervised computational framework [6, 12], GCNG to infer cell-cell interactions from spatial single cell expression data. Our method takes as input both, the location of the cells in the images and the expression of genes in each of these cells. After training it can predict, for any pair of genes, whether they are used in cell-cell interactions for the process being studied. GCNG firstly processed the single cell spatial expression data as one matrix encoding cell locations, and one more matrix encoding gene expression, then feed them into a five-layer graph convolutional neural network to predict gene relationships between cells (Fig. 1A). The core structure of GCN is its graph convolutional layer, which enables it to combine graph structure (cell location) and node information (gene expression in specific cell) as inputs to a neural network. Since the graph structure encodes spatially related cells, GCNs can utilize convolutional layers that underly much of the recent success of neural networks, without directly using image data [29, 31].

**Fig. 1.**
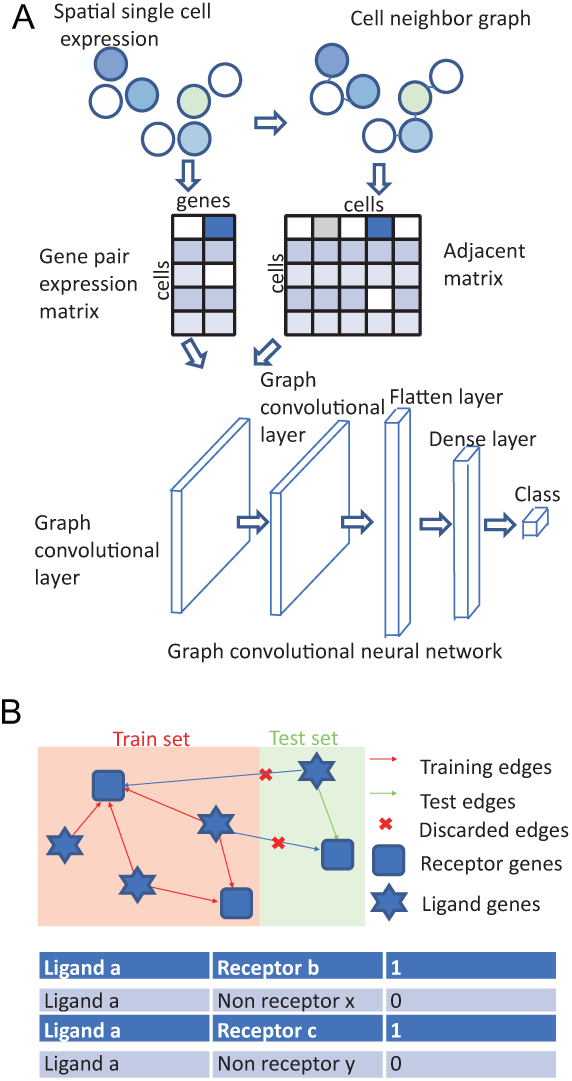
GCNG for extracellular gene relationship inference. (A) GCNG model using spatial single cell expression data. A binary cell adjacent matrix and an expression matrix are extracted from spatial data. After normalization, both matrices are fed into the graph convolutional network. (B) Training and test data separation and generation strategy. The known ligand and receptor genes can form complicated directed networks. For cross validation, all ligand and receptors are separated exclusively as training and test gene sets, and only gene pairs where both genes are in training (test) are used for training (test). To balance the dataset, each positive ligand-receptor (*L*_*a*_, *R*_*b*_) gene pair with label 1 will have a negative pair sample (*L*_*a*_, *R*_*x*_) with label 0 where *R*_*x*_ was randomly selected from all training (test) receptor genes which are not interacting with *L*_*a*_ in training (test).

### 2.1 Dataset used

While several recent methods have been suggested for spatial profiling of transcriptomics data [5, 14, 17, 23, 27] we have focused here on the two methods that provide expression levels for the most number of genes in such experiments. The first is seqFISH+. The seqFISH+ data was downloaded from [8] and included information on the expression of 10,000 genes in 913 cells in the mouse cortex profiled in seven separate fields of view. We normalized the expression data such that expression levels for all genes in each cell sum to the same value as was previously done [8]. The second is a very recent dataset from MERFISH. That data consisted of 12,903 genes in 1,368 cells [30]. Unlike the seqFISH+ data that profiled the expression in the mouse cortex, the MERFISH data is from in vitro cultured cells and so does not include a diverse set of cell types. Still, as the authors of the MERFISH paper observed, even within this population there are difference in spatial expression and so the data can be used to study extracellular gene-gene interaction. See Appendix for complete details on both datasets.

### 2.2 Graph representation for spatial transcriptomics data

We next transformed the spatial cell data to a graph representation. To determine the neighbors of each cell we calculated the Euclidean distance in the image coordinates for all cells, and used a distance threshold to select neighbors. The threshold value was selected so that the average number of neighbors for cells is close to 3, which for the 2D images we used seemed to represent the number of neighbors that were in physical contact with the cell (other threshold choices led to similar results, Tab. S1). Given the set of neighbors we constructed an adjacency matrix of size of 913 × 913, which we term *A*. In other words, for the symmetric network *A, A*_*ij*_ = *A*_*ji*_ = 1 iff *i* and *j* are neighbors and 0 otherwise. Using the adjacency matrix *A* the normalized (symmetric) Laplacian matrix *L*_*N*_ is defined as [21]:

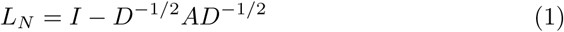

where *D*_*ii*_ = Σ_*j*_ *A*_*ij*_, and *I* is the identity matrix. Other graph representation matrices can also be used in GCNG. One such method which we have also tried first normalizes the adjacency matrix and only after that computes the normalized Laplacian on the normalized adjacency matrix, *L*_*NN*_. See SI for details.

### 2.3 Input construction

GCNG uses two types of input: one encodes the graph structure, as discussed above, while the second encodes node specific values. For this, we first normalized the expression of genes such that the total expression in all cells was the same. Next, we encode the normalized expression of a pair of candidate genes using a matrix of dimension of 913 × 2, (where 913 is the number of cells in our dataset).

### 2.4 Labeled data

GCN requires labeled data for supervised training. While the exact set of signaling interactions between cells in the cortex data we studied is unknown, we used as true interactions a curated list of interacting ligands and receptors [19]. Ligands are proteins that are secreted by cells and they then interact with membrane receptor proteins on neighboring cells to activate signaling pathways within the receiving cell [24]. Several ligand-receptor interactions have been confirmed and here we used a database consisting of 708 ligands, and 691 receptors with 2,557 known interactions [19]. Of these, 309 ligands and 481 involved in 1,056 interactions are also profiled by seqFISH+ and these were used for training and testing.

### 2.5 GCNG network architecture

To construct GCNG we used the python packages of ‘spektral’, ‘Keras’ and ‘Tensorflow’. See Fig. 1A for the architecture of GCNG. GCNG consists of two 32-channel graph convolutional layers, one flatten layer, one 512-dimension dense layer and one Sigmoid function output layer for classification. The graph convolutional layer is defined as:

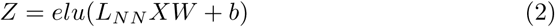

Where *X* is the expression matrix (of dimensions 913 × 2) and *L*_*NN*_ is the 913 × 913 graph matrix discussed above. *W* is a weight matrix of filters (also termed the convolution kernel) with size of 2×32, where 2 corresponds to the two-dimension gene expression of each node, and 32 represents 32 filters or feature maps. *b* is the bias vector term with size of 1 × 32. The ‘*elu*’ (Exponential linear unit) function is defined as:

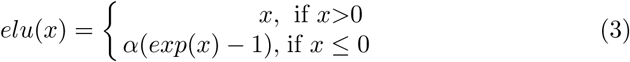

where *α* = 1 by default. Here *Z* represents the embedding vectors of all cell nodes with size of 913 × 32. Note that we are using two convolutional layers here allowing the method to learn indirect graph relationships as well. Since the impact of secreted proteins can be larger than just direct neighbors such an approach allows the method to infer interactions that may be missed by only considering direct neighbors.

The first graph convolutional layer combines the two inputs and converts them to embedding vectors for cell nodes of dimension of 913 × 32. The second graph convolutional layer combines the embedding vector of each cell with the one learned for its direct neighbors. The flatten layer then converts the matrices generated by the second layer to a vector using *ReLU* activation function. Finally a dense layer with one-dimensional output is used to predict the interaction probability based on the *sigmoid* activation function. The activation functions used by the different layers are defined below.

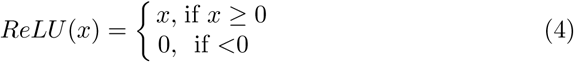

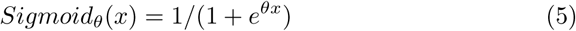

We also tested one and three graph convolutional layers networks, and determined that two layers network led to the best performance.

### 2.6 Train and test strategy

We evaluated GCNG’s performance using ten-fold cross validation. Train and test sets were completely separated to avoid information leakage: For each fold, 90% of ligands and 90% of receptors genes was selected at random for training. All interactions between training proteins were used as a positive set and the negative set was selected from random ligand-receptor pairs that are not known to interact (as mentioned, on average each ligand only interacts with very few receptors so ∼ 99% of random pairs are expected to be negative). Note that pairs for which one of the proteins was part of the training set while another was in the test set were not used and so all proteins in the test set are never seen in training (Fig. 1B). Early stopping through monitoring validation accuracy was used to avoid overfitting, where the last 20% pair samples in training set were selected as validation set, and the patience epoch number was set as half of the total training epoch number. To evaluate models’ performance for ligands, we first calculated the individual Area Under the Receiver Operating Characteristic curve and the Area Under the Precision Recall curve (AUROC/AUPRC) and then combined them for the figures presented.

## 3 Results

### 3.1 GCNG correctly infers ligand-receptor interactions between cells

We first evaluated GCNG’s ability to predict ligand-receptor interactions. For this we used two datasets. The first is seqFISH+ mouse cortex tissue which contains the expression of 10,000 genes in 913 cells. Our labeled set consisted of 1,056 known interactions between 309 ligands and 481 receptors. The second is a MERFISH dataset with 12,903 genes from 1,368 cells, and 841 known interactions between 270 ligands and 376 receptors.

We enforced a strict separation between the training and test sets in the cross validation (Methods, Fig. 1B). Negative pairs were also ligand-receptor and were randomly selected from non-interacting training data genes. To evaluate the performance of our GCNG method we compared it to a number of prior methods that were recently used to predict genes involved in cell-cell interactions from spatial expression data. These include computing the spatial Pearson correlation (PC) between ligand and receptors in neighboring cells (where we used the best neighborhoods for it, see Tab. S1), and Giotto [7] which first calculates a similarity score for all pairs of genes in all pairs of neighboring cell types and then ranks pairs based on their average score. We also compared to two alternative methods that do not use spatial information at all to determine the contribution of neighborhood data. These included Pearson correlation between the expression of ligand and receptors within each cell, and our GCNG method where we only used a diagonal matrix to represent the graph (i.e. each cell is only connected to itself).

Results are presented in Fig. 2 (See Fig. S1 for detailed curves). GCNG reached a mean (median) AUROC/AUPRC of 0.61/0.71 (0.99/1.0). In contrast, for this data spatial PC, Giotto, single cell PC and nonspatial CGNG all performed much worse (mean (median) AUROC/AUPRC of 0.54/0.65 (0.75/0.79), 0.45/0.58 (0.25/0.33), 0.48/0.60 (0.38/0.38) and 0.54,0.64 (0.75/0.79) respectively). In other words, GCNG achieves an improvement of more than 10% for AUROC and more than 20% for AUPRC when compared to all prior methods. In addition to GCNG spatial PC also outperformed the non-spatial PC confirming the importance of spatial information for this task. Testing on the MERFISH data led to similar conclusions. We observed higher mean AUROC/AUPRC values for GCNG when compared to spatial PC (0.64/0.72 for GCNG and 0.59/0.70 for spatial PC). See Tab. S2 for complete results for MERFISH. While GCNG improvement over spatial PC for MERFISH is lower than the improvement observed for seqFISH+, this can be attributed to the type of data analyzed. As noted in Methods, the MERFISH data is less ideal for cell-cell interactions analysis since it only profiles a single cell type (human osteosarcoma (U-2 OS) cells). Given this, we have focused on seqFISH+ for subsequent analysis.

**Fig. 2.**
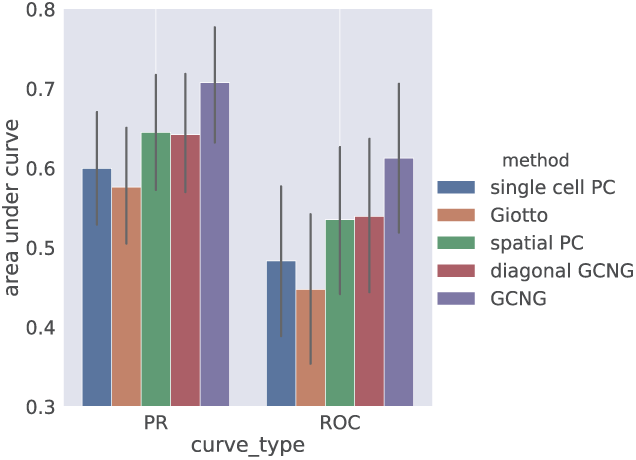
GCNG correctly infers ligand-receptor interaction between neighboring cells. Overall AUROC and AUPRC for GCNG, spatial Pearson correlation, within single cell Pearson correlation, Giotto and GCNG model with a diagonal adjacent matrix without spatial information.

We next selected the top 3000 highly expressed genes for all cells in the seqFISH+ data, and used GCNG trained on the entire ligand-receptor database to predict scores for all possible edges among genes. Manual inspection of the top 10 predicted pairs shows that four of them are supported by recent studies [1, 11, 10, 16] (Tab. 1). For example, the top pair “apt1a1-itgb5”, contains two genes that are known to be involved in membrane interaction functions [16].

**Table 1.**
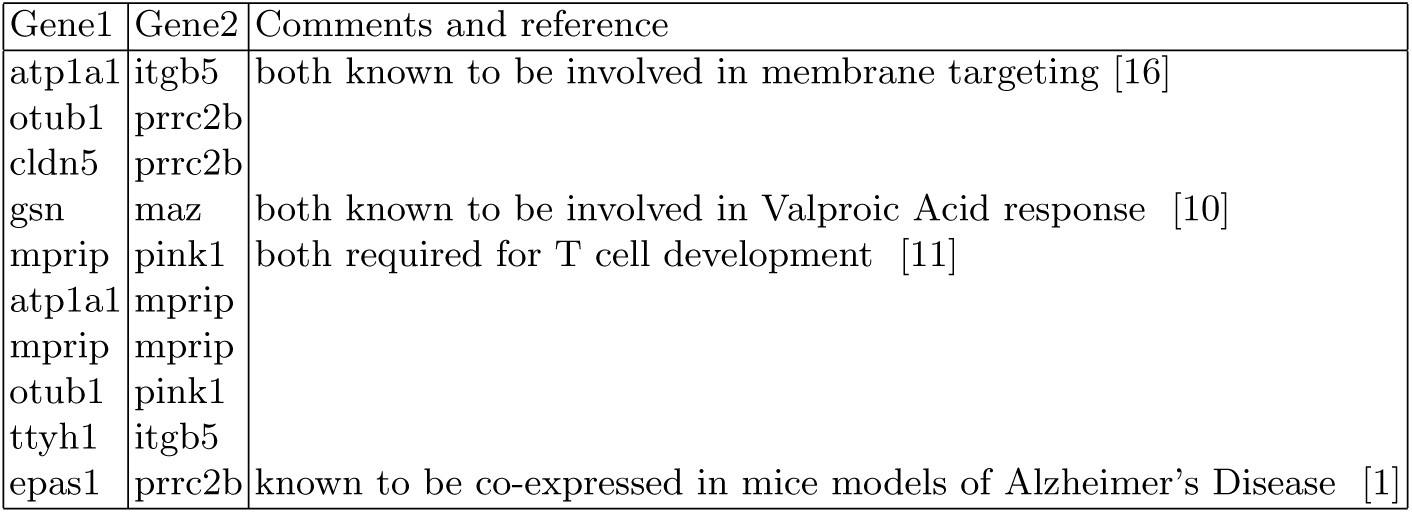
Top predicted gene pairs from highly expression genes. Four of them are supported by recent studies.

### 3.2 Analysis of co-expression patterns identified by GCNG

To further explore the predictions of our GCNG and to map them back to the original spatial representation we looked at some of the top correctly predicted pairs. For each such pair (a ligand and receptor predicted to interact) we projected their spatial expression on the cell distribution figures. Fig 3 presents such projection for two ligands (col4a1 and lamc1) with their positive and negative receptor partners (Fig. 3A, B for col4a1, and C, D for lamc1) (See Fig. S2 for plots of all fields of view). Here, cells are defined to ‘highly expressing’ a gene if the expression of the gene in that cell is in the top 100 expression levels for that gene. For the positive col4a1-cd93 pair, cells highly expressing col4a1 and cd93 are both concentrated in the 1st and 5th fields. In contrast, for the negative col4a1-hrh3 pair, cells highly expressing hrh3 do not seem to reside next to cells expressing col4a1. Similar pattern comparison can also be observed for ligand lamc1 with positive (itgb1) and negative receptor (lyve1) (Fig. 3C, D). The ability of GCNG to predict such interactions based on a subset of the data highlights the usefulness of this approach compared to global analysis methods including PC. Cell type plots (Fig. S3) indicate that correctly predicted pairs can be found in both, neighboring cells from the same type and cells from different types. These results indicate that the GCNG method can generalize well and can be used to correctly identify several different types of interactions.

**Fig. 3.**
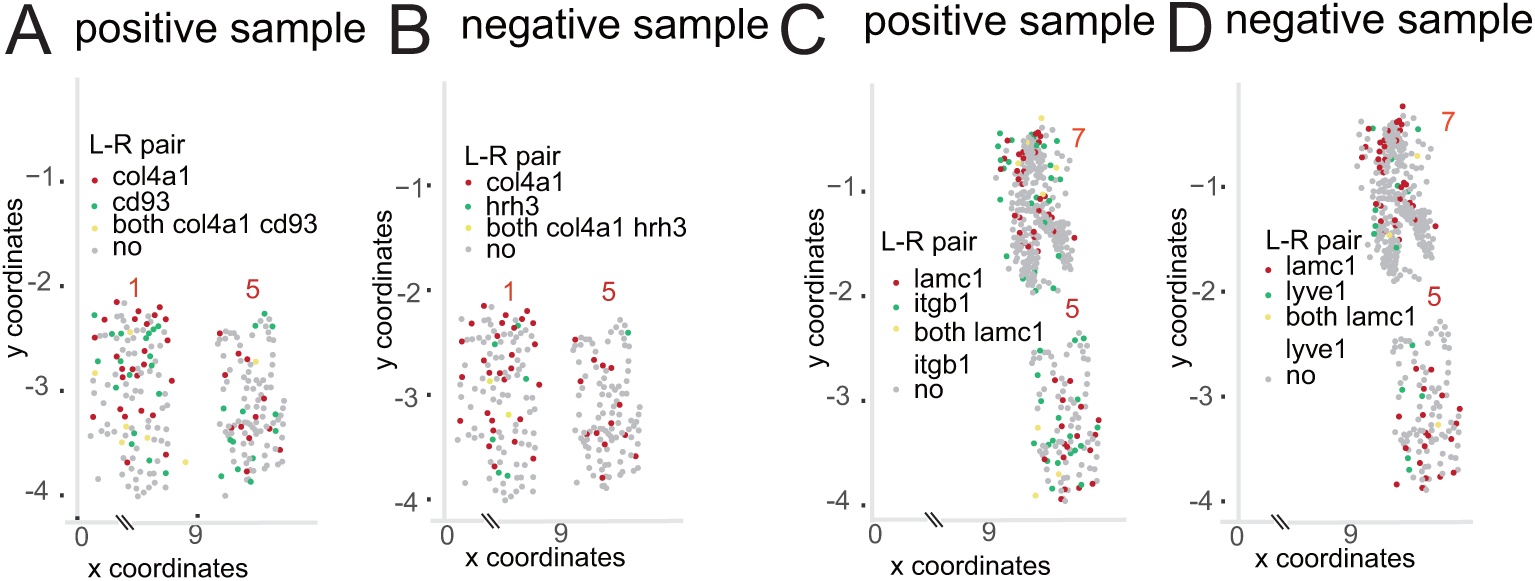
Spatial expression patterns for selected ligand-receptor pairs. (A, B) Spatial expression distribution of correctly predicted positive (cd93) and negative (hrh3) receptors for ligand col4a1. 2 of the 7 fields of view (FOV) profiled are shown (see Appendix for all FOVs). As can be seen, the correctly predicted pairs are indeed much better spatially correlatd than the negative pair. (C, D) Spatial expression distribution of correctly predicted positive (itgb1) and negative (lyve1) receptors for ligand lamc1. Cells highly expressing lamc1 and itgb1 are both concentrated in the 5th and 7th fields as shown.

### 3.3 Inferring causal interactions

While correlation-based methods can be used to identify gene co-expression interactions and networks [3, 13], these methods cannot be used to infer causality since their outcome is symmetric. Causality information may be trivial for ligand-receptors, since the direction for such pair is known. However, for other interacting genes across cells the direction is often not clear. Thus, a method that can infer both interactions and directionality may be beneficial for studying spatial transcriptomics data. Unlike prior unsupervised methods, our supervised framework can be trained to identify causal interactions if training data exists. We thus trained a GCNG on a subset of known causal pairs (ligand and receptors) and then used it to predict directionality for other pairs. To generate train and test data for this, for each known ligand-receptor (*L*_*a*_, *R*_*b*_) gene pair we introduce a negative sample (*R*_*b*_, *L*_*a*_) with label 0. The same ten-fold cross validation strategy is used to evaluate GCNG’s performance here. Results are presented in Fig. 4. As can be seen GCNG performs well on this task with mean (median) AUROC of 0.60 (0.99) and AUPRC of 0.70 (1.0) respectively. Thus, for top predicted pairs, the direction predictions of GCNG can be used to further assign causality.

**Fig. 4.**
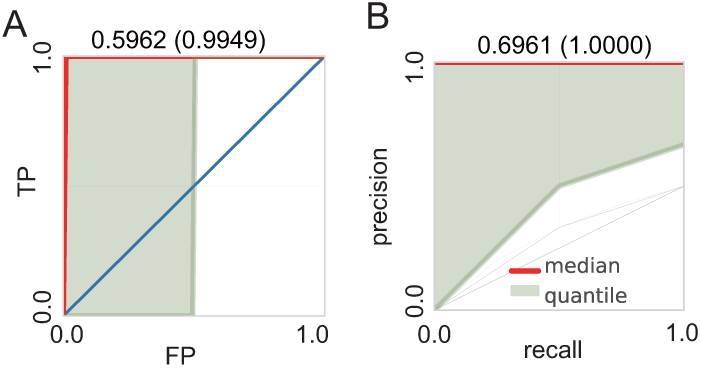
GCNG can infer direction of ligand-receptor interactions between cells. (A, B) AUROC and AUPRC of GCNG on the direction prediction task. Here each gray line represents one ligand, red line represents the median curve, and the light green part represents the region between 40-60 quantile. Mean and median of area under the curves are shown on top of each panel.

### 3.4 Functional gene assignment

We next tested whether GCNG can be used for applications that utilize predicted interactions as features for down stream analysis. Specifically, we tested whether the outcomes of GCNG can be used as features for assigning function to genes. A popular method for such assignment is Guilt By Association (GBA) [18]. In GBA candidate gene association with known genes is calculated, the total value of which is then used as the final score for this candidate. For this as an alternative to GBA, we trained GCNG to distinguish the spatial expression of pairs of genes within the same function (positive set) from pairs where one gene is associated with that function and the other is not (negative). We focused on functions related to cell-cell communication. In this analysis we used the GSEA sets [25] for integrin cell surface interactions, cell surface interactions at the vascular wall and cell-cell communication, which consist of 70, 77 and 79 genes in the seqFISH+ data, respectively. Performance was evaluated using five-fold cross validation. Since in function assignment tasks validation experiments are usually limited to the top few genes we focused the evaluation on the top 20% predictions based on the scores of GBA and GCNG. Results are presented in Fig. 5 and indicate that for communication related functions using spatial information can improve functional gene assignment.

**Fig. 5.**
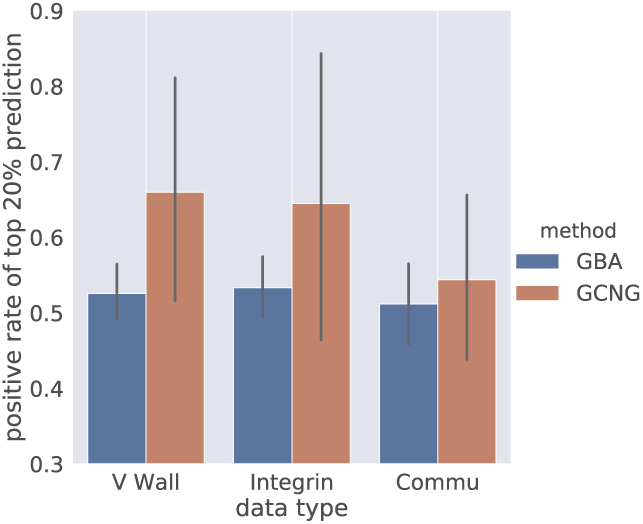
Function assignment. Fraction of correctly assigned genes in top 20% of predictions for GCNG and GBA for ‘integrin cell surface interactions’, ‘cell surface interactions at the vascular wall’ and ‘cell-cell communication’ functions respectively.

## 4 Discussion

Gene expression data has been extensively and successfully used to infer interaction between genes, gene regulation and temporal and causal effects [4, 9, 22]. With the recent advances in spatial transcriptomics such data can now be used to infer pairs of genes involved in cell-cell communication. However, directly converting methods used to infer intra-cellular interactions to methods for inferring extra-cellular interactions is not trivial. The spatial data tends to be very sparse, contains several different cell types and requires specific decisions about the neighborhoods to consider.

To overcome these problems we developed a supervised GCN approach for the analysis of spatial transcriptomics data. GCN has been used in the past for computational biology, though prior applications did not focus on cells but rather on intracellular pathways [32]. In addition, prior work utilized known interaction to define the graph structure. In contrast, our method can work with a general spatial image for which the specific interactions between cells are not known. The method generates a neighborhood graph based on distances between cells and uses it, together with the pairwise expression values for genes to predict interactions between and across cells.

Application of our CGNG method to datasets that provide the highest coverage of genes shows that it can successfully identify known ligand-receptor pairs and that it is much more accurate when compared to prior methods proposed for this or to methods that do not utilize the spatial information. Visualization of some resulting predictions highlights the ability of GCNG to focus on a relevant subset of locations rather than on global correlation. The output of GCNG can also be used as features for down-stream analysis including for methods for gene function assignment and methods for learning interaction networks.

While successful, there are several ways in which GCNG can be improved. First, the choice of the number of convolutional layers to use (which relates to the assumption about the propagation distance of secreted proteins) needs to be better handled to fit the needs of individual datasets. Another important direction is to focus more on specific cell types rather than on the overall interactions. We expect to address these and other issues in future work.

GCNG is implemented in Python and both data and an open source version of the software are available from the supporting website.

## References

1. Bouter, Y., Kacprowski, T., Weissmann, R., Dietrich, K., Borgers, H., Brauß, A., Sperling, C., Wirths, O., Albrecht, M., Jensen, L.R., et al.: Deciphering the molecular profile of plaques, memory decline and neuron loss in two mouse models for alzheimer’s disease by deep sequencing. Frontiers in aging neuroscience 6, 75 (2014)

2. Bruna, J., Zaremba, W., Szlam, A., LeCun, Y.: Spectral networks and locally connected networks on graphs. arXiv preprint 1312.6203 (2013)

3. Care, M.A., Westhead, D.R., Tooze, R.M.: Parsimonious gene correlation network analysis (pgcna): a tool to define modular gene co-expression for refined molecular stratification in cancer. NPJ systems biology and applications 5(1), 13 (2019)

4. Chan, T.E., Stumpf, M.P., Babtie, A.C.: Gene regulatory network inference from single-cell data using multivariate information measures. Cell systems 5(3), 251–267 (2017)

5. Codeluppi, S., Borm, L.E., Zeisel, A., La Manno, G., van Lunteren, J.A., Svensson, C.I., Linnarsson, S.: Spatial organization of the somatosensory cortex revealed by osmfish. Nature methods 15(11), 932 (2018)

6. Defferrard, M., Bresson, X., Vandergheynst, P.: Convolutional neural networks on graphs with fast localized spectral filtering. In: Advances in neural information processing systems. pp. 3844–3852 (2016)

7. Dries, R., Zhu, Q., Eng, C.H.L., Sarkar, A., Bao, F., George, R.E., Pierson, N., Cai, L., Yuan, G.C.: Giotto, a pipeline for integrative analysis and visualization of single-cell spatial transcriptomic data (2019)

8. Eng, C.H.L., Lawson, M., Zhu, Q., Dries, R., Koulena, N., Takei, Y., Yun, J., Cronin, C., Karp, C., Yuan, G.C., et al.: Transcriptome-scale super-resolved imaging in tissues by rna seqfish+. Nature 568(7751), 235 (2019)

9. Hill, S.M., Heiser, L.M., Cokelaer, T., Unger, M., Nesser, N.K., Carlin, D.E., Zhang, Y., Sokolov, A., Paull, E.O., Wong, C.K., et al.: Inferring causal molecular networks: empirical assessment through a community-based effort. Nature methods 13(4), 310 (2016)

10. Jergil, M., Kultima, K., Gustafson, A.L., Dencker, L., Stigson, M.: Valproic acid– induced deregulation in vitro of genes associated in vivo with neural tube defects. Toxicological sciences 108(1), 132–148 (2009)

11. Kakoola, D.N., Curcio-Brint, A., Lenchik, N.I., Gerling, I.C.: Molecular pathway alterations in cd4 t-cells of nonobese diabetic (nod) mice in the preinsulitis phase of autoimmune diabetes. Results in immunology 4, 30–45 (2014)

12. Kipf, T.N., Welling, M.: Semi-supervised classification with graph convolutional networks. arXiv preprint 1609.02907 (2016)

13. Langfelder, P., Horvath, S.: Wgcna: an r package for weighted correlation network analysis. BMC bioinformatics 9(1), 559 (2008)

14. Lee, J.H., Daugharthy, E.R., Scheiman, J., Kalhor, R., Yang, J.L., Ferrante, T.C., Terry, R., Jeanty, S.S., Li, C., Amamoto, R., et al.: Highly multiplexed subcellular rna sequencing in situ. Science 343(6177), 1360–1363 (2014)

15. Lin, C., Jain, S., Kim, H., Bar-Joseph, Z.: Using neural networks for reducing the dimensions of single-cell rna-seq data. Nucleic acids research 45(17), e156–e156 (2017)

16. Misselwitz, B., Dilling, S., Vonaesch, P., Sacher, R., Snijder, B., Schlumberger, M., Rout, S., Stark, M., Von Mering, C., Pelkmans, L., et al.: Rnai screen of salmonella invasion shows role of copi in membrane targeting of cholesterol and cdc42. Molecular systems biology 7(1) (2011)

17. Moffitt, J.R., Bambah-Mukku, D., Eichhorn, S.W., Vaughn, E., Shekhar, K., Perez, J.D., Rubinstein, N.D., Hao, J., Regev, A., Dulac, C., et al.: Molecular, spatial, and functional single-cell profiling of the hypothalamic preoptic region. Science 362(6416), eaau5324 (2018)

18. Oliver, S.: Proteomics: guilt-by-association goes global. Nature 403(6770), 601 (2000)

19. Ramilowski, J.A., Goldberg, T., Harshbarger, J., Kloppmann, E., Lizio, M., Satagopam, V.P., Itoh, M., Kawaji, H., Carninci, P., Rost, B., et al.: A draft network of ligand–receptor-mediated multicellular signalling in human. Nature communications 6, 7866 (2015)

20. Sanderson, C.M.: A new way to explore the world of extracellular protein interactions. Genome research 18(4), 517–520 (2008)

21. Shuman, D.I., Narang, S.K., Frossard, P., Ortega, A., Vandergheynst, P.: The emerging field of signal processing on graphs: Extending high-dimensional data analysis to networks and other irregular domains. IEEE signal processing magazine 30(3), 83–98 (2013)

22. Song, L., Langfelder, P., Horvath, S.: Comparison of co-expression measures: mutual information, correlation, and model based indices. BMC bioinformatics 13(1), 328 (2012)

23. Ståhl, P.L., Salmén, F., Vickovic, S., Lundmark, A., Navarro, J.F., Magnusson, J., Giacomello, S., Asp, M., Westholm, J.O., Huss, M., et al.: Visualization and analysis of gene expression in tissue sections by spatial transcriptomics. Science 353(6294), 78–82 (2016)

24. Stassen, F.L.: Molecular foundations of drug-receptor interaction. cambridge university press, cambridge, london, new york, new rochelle, melbourne, sydney 1987. 381 pp. Journal of Molecular Recognition 1(3), ii–ii (1988)

25. Subramanian, A., Tamayo, P., Mootha, V.K., Mukherjee, S., Ebert, B.L., Gillette, M.A., Paulovich, A., Pomeroy, S.L., Golub, T.R., Lander, E.S., et al.: Gene set enrichment analysis: a knowledge-based approach for interpreting genome-wide expression profiles. Proceedings of the National Academy of Sciences 102(43), 15545–15550 (2005)

26. Van Dijk, D., Sharma, R., Nainys, J., Yim, K., Kathail, P., Carr, A.J., Burdziak, C., Moon, K.R., Chaffer, C.L., Pattabiraman, D., et al.: Recovering gene interactions from single-cell data using data diffusion. Cell 174(3), 716–729 (2018)

27. Wang, X., Allen, W.E., Wright, M.A., Sylwestrak, E.L., Samusik, N., Vesuna, S., Evans, K., Liu, C., Ramakrishnan, C., Liu, J., et al.: Three-dimensional intact-tissue sequencing of single-cell transcriptional states. Science 361(6400), eaat5691 (2018)

28. Wei, Z., Li, H.: A markov random field model for network-based analysis of genomic data. Bioinformatics 23(12), 1537–1544 (2007)

29. Wu, Z., Pan, S., Chen, F., Long, G., Zhang, C., Yu, P.S.: A comprehensive survey on graph neural networks. arXiv preprint 1901.00596 (2019)

30. Xia, C., Fan, J., Emanuel, G., Hao, J., Zhuang, X.: Spatial transcriptome profiling by merfish reveals subcellular rna compartmentalization and cell cycle-dependent gene expression. Proceedings of the National Academy of Sciences 116(39), 19490–19499 (2019)

31. Zhou, J., Cui, G., Zhang, Z., Yang, C., Liu, Z., Sun, M.: Graph neural networks: A review of methods and applications. arXiv preprint 1812.08434 (2018)

32. Zitnik, M., Agrawal, M., Leskovec, J.: Modeling polypharmacy side effects with graph convolutional networks. Bioinformatics 34(13), i457–i466 (2018)

