## Supplement for "GCNG: Graph convolutional networks for inferring cell-cell interactions"

**Methods:**

**Expression normalization**

The counts in seqFISH+ data were processed with the following rule:

$$expresion_{ij} = \frac{count_{ij}}{\sum_j count_{ij}} \times 10{,}000$$

Where *i* represents cell *i*, and *j* represents gene *j*.

**Graph matrices**

The normalized (symmetric) Laplacian matrix $\boldsymbol{L_N}$ was generated by setting:

$L_N = I - D^{-1/2}AD^{-1/2}$, Where $D_{ii} = \sum_j A_{ij}$, *I* is the identity matrix. (1)

$\boldsymbol{L_N}$ can be the operator approximation of 1st order Chebyshev polynomials approximation (CPA) for original spectral convolutions on graphs [1], with the assumption that its zero order is the opposite number of its 1st order coefficient [2]. Specifically, the spectral convolutions on graphs is defined as $g_\theta * x = U g_\theta(\Lambda) U^T x$, where *U* is matrix of eigenvectors of normalized Laplacian matrix $\boldsymbol{L_N}$ from *A* (Eq. 1), *Λ* is diagonal matrix of eigenvalues of $\boldsymbol{L_N}$, $U^T x$ is the graph Fourier transform of $x$, $g_\theta$ is a function of *Λ*. The K$^{th}$ order CPA $g_{\theta'}(\Lambda) \approx \sum_{k=0}^{K} \theta'_k T_k(\tilde{\Lambda})$ was then used to approximate $g_\theta(\Lambda)$, where $\tilde{\Lambda} = 2\Lambda / \lambda_{max} - I_N$, $\lambda_{max}$ is the largest entry of $\Lambda$, $T_k(x) = 2xT_{k-1}(x) - T_{k-2}(x)$, $T_0(x) = 1$ and $T_1(x) = x$. The 1st order CPA can be approximated as $\theta_0 x -$

$\theta_1 D^{-1/2} A D^{-1/2} x$ with $\lambda_{max} \approx 2$. By further assumption of $\theta = \theta_0 = \theta_1$, the final approximation becomes:

$$g_\theta * x \approx \theta(I - D^{-1/2} A D^{-1/2})x \qquad (2)$$

We also tried two alternative graph matrix approximation. a) The normalized Laplacian matrix can be generated from a normalized adjacent matrix, $\boldsymbol{A_N}$ instead of $\boldsymbol{A}$:

$$A_N = D^{-1/2} A D^{-1/2}, \text{ Where } D_{ii} = \sum_j A_{ij}. \qquad (3)$$

Then the normalized (symmetric) Laplacian matrix $\boldsymbol{L_{NN}}$ from $\boldsymbol{A_N}$ was generated as the graph matrix:

$$L_{NN} = I - D_N^{-1/2} A_N D_N^{-1/2}, \text{ Where } D_{N_{ii}} = \sum_j A_{N_{ij}}. \qquad (4)$$

b) Following the same assumption of $\theta = \theta_0 = -\theta_1$ in ref [2], and renormalized trick of $I + D^{-1/2} A D^{-1/2} \to D'^{-1/2} A' D'^{-1/2}$, the convolutional approximation becomes $g_\theta * x \approx \theta D'^{-1/2} A' D'^{-1/2} x$, Where $D'_{ii} = \sum_j A'_{ij}$, and $A' = A + I$. Then, graph matrix approximation can be generated by setting, $G = D'^{-1/2} A' D'^{-1/2}$, Where $D'_{ii} = \sum_j A'_{ij}$, and $A' = A + I$. The performance of all the three graph matrices were much better than other methods in the ligand-receptor interaction task (See **Tab. S1** for details). Since the graph matrix $\boldsymbol{L_{NN}}$ also performed well in other tasks, it was the one used for all tasks across the paper.

**Figures:**

**Fig. S1 Detailed AUROC and AUPRC for Fig. 2**

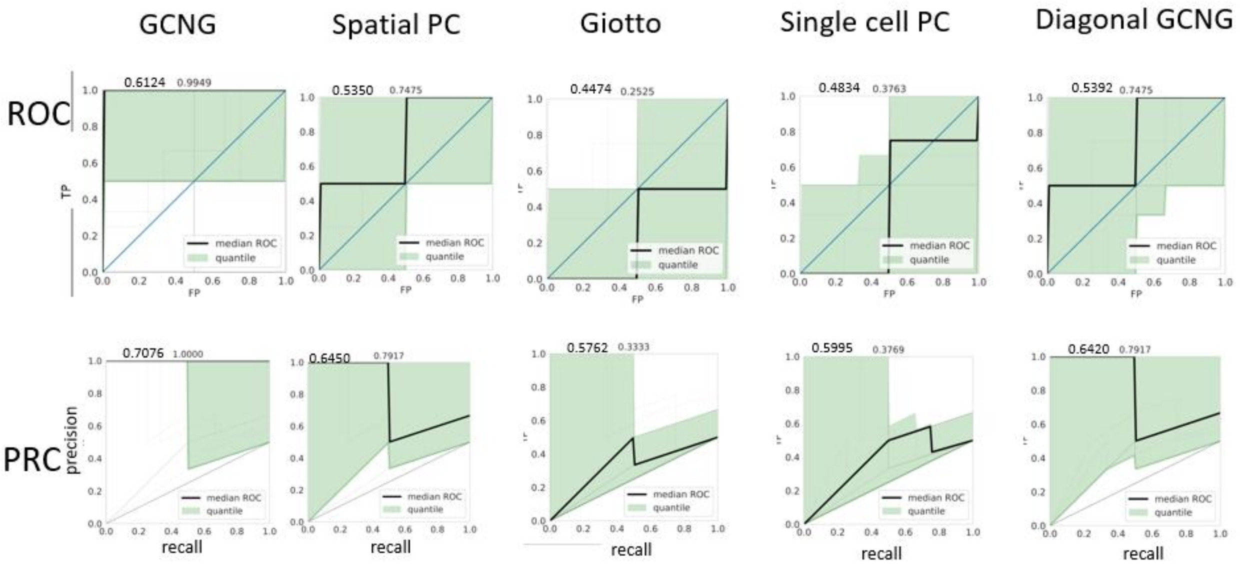


Here each gray line represents one ligand, black line represents the median curve, and the light green part represents the region between 40-60 quantile. Mean and median of area under the curves are shown in top of each panel.

**Fig. S2 Whole plots of typical gene pair's spatial expression pattern for Fig. 3**

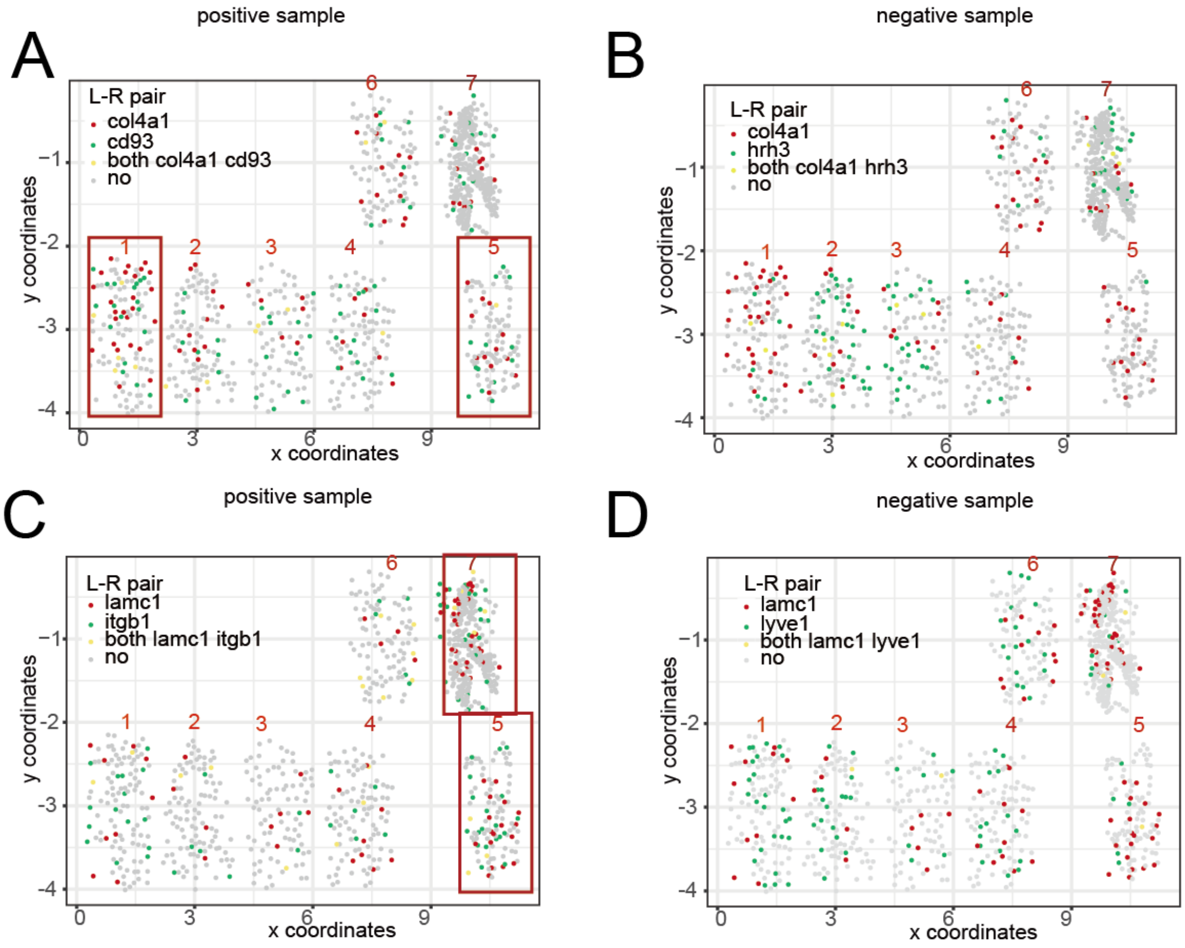


**(A, B)** The spatial expression distribution of correctly predicted positive (cd93) and negative (hrh3) samples for ligand col4a1. Cells highly expressing col4a1 (red) and cd93 (green) are both concentrated in the 1st and 5th fields as shown in the red boxes (Fig. 3). **(C, D)** The spatial expression distribution of correctly predicted positive (itgb1) and negative (lyve1) samples for ligand lamc1. Cells highly expressing lamc1 and itgb1 are both concentrated in the 5th and 7th fields as shown in the red boxes.

**Fig. S3 Cell type spatial distribution**

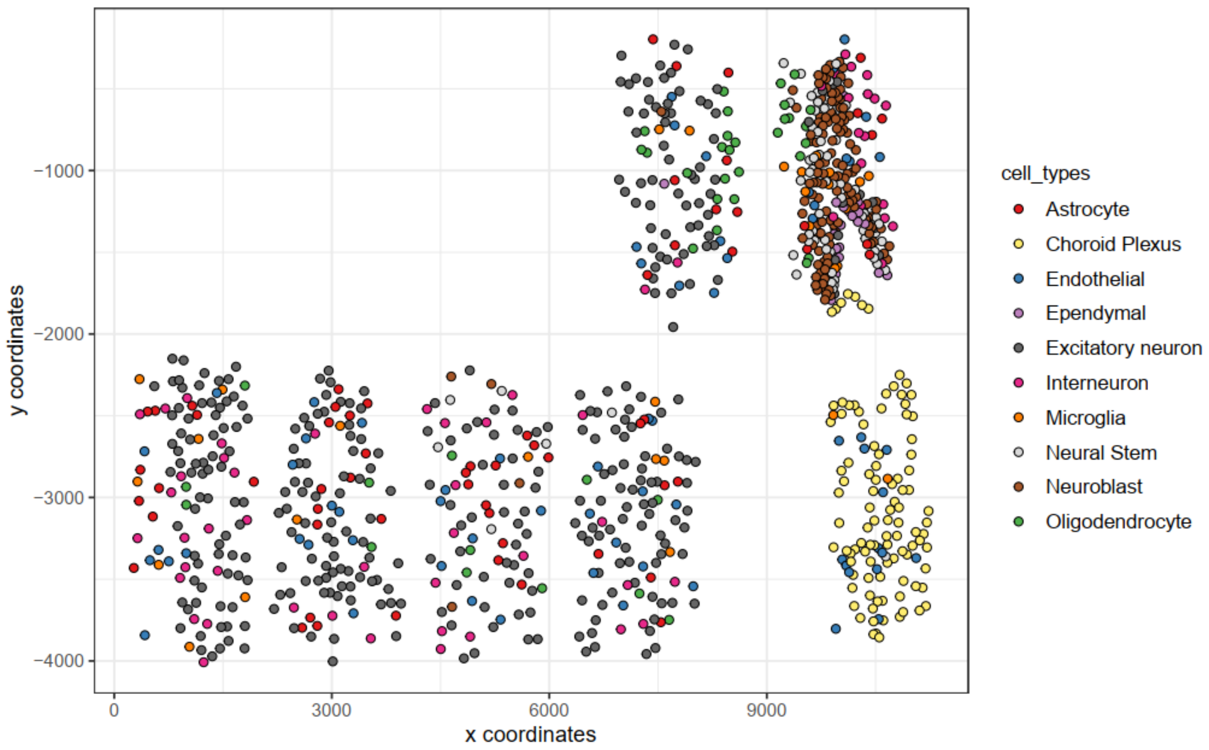


**Tables:**

**Tab. S1**

**Performance on ligand-receptor interaction prediction task of GCNG with three different input graph matrices and Spatial PC using a series of distance thresholds, based on seqFISH+ data**

| Distance Threshold | | 0 (diagonal) | 100 | 140 | 180 | 220 | 260 |
|---|---|---|---|---|---|---|---|
| Average neighbor count | | 0 | 1.2 | 2.5 | 4.0 | 5.8 | 7.7 |
| $\underline{D'^{-1/2}A'D'^{-1/2}}$ | Average AUROC | 0.5392 | 0.5006 | 0.6256 | 0.6462 | 0.6390 | 0.5446 |
| | Average AUPRC | 0.6420 | 0.6128 | 0.7114 | 0.7282 | 0.7194 | 0.6519 |
| Laplacian matrix from Normalized adjacent matrix | Average AUROC | 0.5392 | 0.5978 | 0.6124 | 0.5131 | 0.5249 | 0.5165 |
| | Average AUPRC | 0.6420 | 0.6963 | 0.7076 | 0.6283 | 0.6322 | 0.6303 |
| Laplacian matrix from adjacent matrix | Average AUROC | 0.5392 | 0.5616 | 0.5797 | 0.5147 | 0.4711 | 0.4885 |
| | Average AUPRC | 0.6420 | 0.6697 | 0.6811 | 0.6260 | 0.5956 | 0.6143 |
| Spatial PC | Average AUROC | | 0.4193 | 0.4697 | 0.5350 | 0.5103 | 0.5053 |
| | Average AUPRC | | 0.5610 | 0.6027 | 0.6450 | 0.6363 | 0.6224 |

The red numbers represent the best performance of the methods respectively.

**Tab. S2**

**Performance on ligand-receptor interaction prediction task of GCNG with Laplacian matrix from normalized adjacent matrix and Spatial PC using a series of distance thresholds, based on MERFISH data**

| Distance threshold | | 0 | 110 | 130 | 150 | 170 | 200 | 240 | 280 |
|---|---|---|---|---|---|---|---|---|---|
| Average neighbor count | | 0 | 0.59 | 1.04 | 1.52 | 2.05 | 2.95 | 4.25 | 5.73 |
| Spatial PC | Average AUROC | 0.5886 | 0.5397 | 0.5431 | 0.5301 | 0.5934 | 0.5903 | 0.5485 | 0.5583 |
| | Average AUPRC | 0.6806 | 0.6602 | 0.6553 | 0.6467 | 0.6965 | 0.6913 | 0.6583 | 0.6647 |
| GCNG | Average AUROC | 0.5780 | 0.5867 | 0.6082 | 0.6362 | 0.5546 | 0.5312 | 0.5139 | 0.5758 |
| | Average AUPRC | 0.6779 | 0.6883 | 0.7050 | 0.7218 | 0.6681 | 0.6441 | 0.6328 | 0.6833 |

Giotto is not used here because all cells are the same cell type, Human Bone Osteosarcoma Epithelial Cells (U2OS Line). The purple values of Spatial PC with 0 threshold represent the single cell PC results. The values of GCNG with 0 threshold represent the diagonal GCNG results.

**Reference:**

1. Defferrard, M., Bresson, X., Vandergheynst, P.: Convolutional neural networks on graphs with fast localized spectral filtering. In: Advances in neural information processing systems. pp. 3844{3852 (2016)

2. Kipf, T.N., Welling, M.: Semi-supervised classification with graph convolutional networks. arXiv preprint arXiv:1609.02907 (2016)
